# Meta-analysis of gut microbiome in largemouth bass (Micropterus salmoides) under different aquaculture systems

**DOI:** 10.64898/2026.09.13.751318

**Authors:** Xiangnan Liu, Liang Zhong, Weiwei Zeng, Wenlong Cai

## Abstract

The teleost gastrointestinal tract hosts metabolically active microbial assemblages that play critical roles in nutrient use, immune regulation, physiology, behavior, growth, reproduction, and health status. However, the extent to which aquaculture production systems restructure these communities remains poorly defined. In this study, metagenomic sequencing was applied to characterize gut microbiota in largemouth bass (*Micropterus salmoides*) reared under four representative farming conditions: industrial aquaculture (IA), captive pond-based farming (CA), ordinary pond aquaculture with healthy fish (PH), and ordinary pond aquaculture with diseased fish (PD). Results revealed that the four groups exhibited pronounced differences in microbial diversity, taxonomic composition, functional capacity, carbohydrate-active enzyme profiles, and antibiotic resistance genes (ARGs). Notably, bacterial taxa accounted for more than 95% of the gut microbial communities across all groups. IA fish showed greater microbial richness and diversity than CA and PH fish, consistent with a more complex and potentially more stable intestinal ecosystem under industrialized rearing conditions. Taxonomic profiling at both the phylum and genus levels identified production system-specific microbial signatures. In particular, opportunistic pathogens, including *Aeromonas veronii* and *Aeromonas hydrophila*, were disproportionately enriched in CA fish, suggesting a microbiota profile associated with increased disease vulnerability. Functional prediction analysis further indicated distinct metabolic states among groups, with purine and fatty acid metabolism enriched in PD fish, potentially reflecting disease-associated disruption of host-microbiota interactions. Carbohydrate-active enzyme analysis revealed substantial variation in microbial carbohydrate-processing potential, whereas ARG profiling indicated that farming conditions were linked to distinct gut resistome patterns. Collectively, these findings identify aquaculture regimes as a key ecological driver of intestinal microbiome structure in largemouth bass and provide a mechanistic basis for optimizing microbiome-informed health management in aquaculture systems.

## 1. Introduction

Largemouth bass (*Micropterus salmoides*), a member of the order Perciformes and family Centrarchidae, is indigenous to the southeastern United States, northeastern Mexico, and southeastern Canada (Xie et al. 2023; Zhao, Yang, et al. 2023; Zhao, Liu, et al. 2023; Bai et al. 2008). Given its rapid growth, broad tolerance to diverse culture conditions, relatively short production cycle, and favorable flesh quality, this species has become highly favored among consumers (Sun et al. 2021). Following the introduction of the northern strain from Taiwan into mainland China in 1983, advances in formulated feed and domestication of feeding behavior accelerated expansion of largemouth bass aquaculture (Wang et al. 2019). As production has intensified, research has increasingly focused on growth optimization, immune competence, and disease control in this species (Liao et al. 2023; Li et al. 2024; Liu et al. 2025). Within this context, the intestinal microbiome provides a biological link between rearing conditions and host performance. In addition to its well-established roles in digestion, nutrient absorption, and immunomodulation, gut microbial communities exhibit considerable plasticity and respond sensitively to environmental conditions, dietary composition, and management practices in aquaculture systems (Liu et al. 2021; Austin 2002; Rawls et al. 2006).

Industrial aquaculture (IA), cage aquaculture (CA), and pond aquaculture (PA) impose distinct ecological pressures through differences in water quality, stocking density, feed input, and microbial exposure, each of which can reshape gut microbial composition and diversity (Xiong, Nie, and Chen 2019; Tremaroli and Bäckhed 2012; Zhao et al. 2018; Ray et al. 2019). Despite increasing recognition that intestinal microbiota influence fish health and productivity, comparative metagenomic evidence across major farming systems remains limited for largemouth bass. In particular, the extent to which production-system-dependent microbial shifts alter functional potential, pathogen-associated taxa, carbohydrate metabolism, and antibiotic resistance profiles remains poorly resolved. This research gap limits development of microbiome-guided strategies for disease prevention and sustainable intensification of largemouth bass aquaculture.

This study applied metagenomic sequencing to comprehensively characterize the gut microbiota of largemouth bass reared under different aquaculture regimes. The analysis was designed to identify production-system-specific microbial signatures, characterize functional variation among farming conditions, and establish a microbiological basis for refining aquaculture management through targeted regulation of intestinal microbial communities.

## 2. Materials and Methods

### 2.1. Fish and sampling

Largemouth bass (*Micropterus salmoides*) fry originating from a single batch were randomly allocated to three aquaculture systems, with three replicate units per system: pond captive mode (CA group), industrial aquaculture mode using a land-based recirculating aquaculture system (IA group), and ordinary pond culture mode (PH group) (Figure 1). All fish received the same commercial diet (Zhejiang Xinxin Feed, Zhejiang, China) at 2% of body weight. During culture, dissolved oxygen was maintained above 5.0 mg/L, pH was maintained at 6.5–7.5, and total ammonia nitrogen did not exceed 1 mg/L. Fish were cultured from August to November 2024, during which water temperature ranged from 19.4 to 30.5 °C. After approximately three months of cultivation, fish reached an average body weight of 450 ± 45 g, and intestinal samples were collected. An outbreak of largemouth bass virus (LMBV) occurred in one pond culture during the experiment. Fish from this pond were sampled before chemical treatment and assigned to a fourth group, defined as pond aquaculture with diseased fish (PD group).

**Figure 1.**
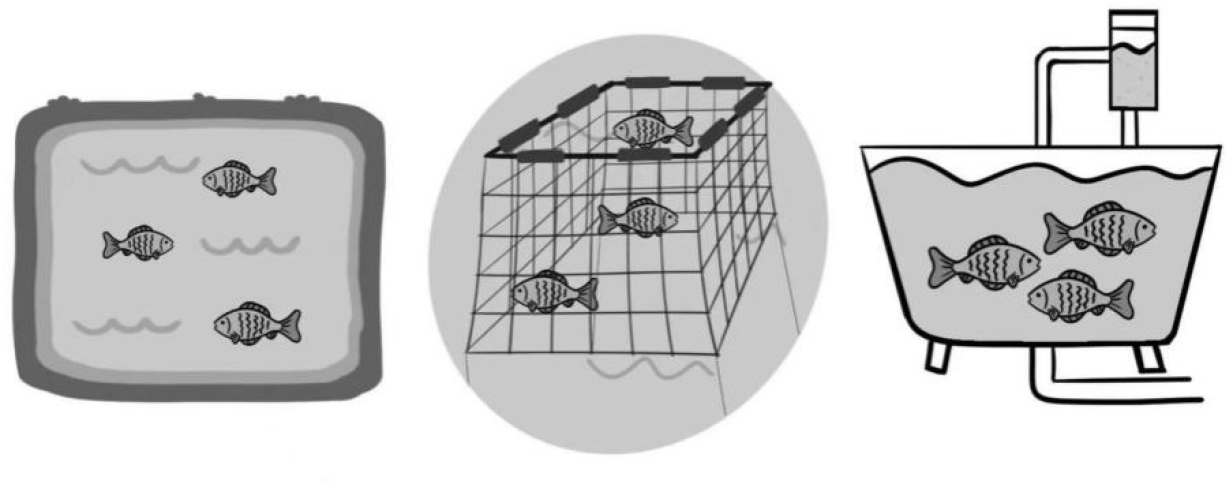
Representative images of the three culture systems. Ordinary pond culture mode (PH group), pond captive mode (CA group), and land-based recirculating industrial aquaculture mode (IA group).

Before sampling, all fish were fasted for 12 h and then enthanized with an overdose of MS-222. Five intestinal samples were collected from each group, and each sample consisted of pooled intestinal contents from three fish. Under sterile conditions, the ventral body wall was incised to expose the peritoneal cavity. The entire intestine was removed, rinsed repeatedly with 0.65% sterile saline, and processed to collect intestinal contents into sterile Eppendorf tubes. Samples were stored at −80 °C until subsequent analysis.

### 2.2. DNA extraction, library construction, and metagenomic sequencing

Total genomic DNA was extracted from intestinal content samples using a QIAamp PowerFecal Pro DNA Kit (Qiagen, Hilden, Germany) according to the protocols supplied by the manufacturer. DNA concentration and purity were determined using a TBS-380 fluorometer and NanoDrop 2000 spectrophotometer, respectively, and DNA integrity was assessed by 1% agarose gel electrophoresis. Extracted DNA was sheared to an average fragment size of approximately 350 bp using a Covaris M220 ultrasonicator (Gene Company Limited, China), followed by paired-end library preparation. Sequencing libraries were generated with a NEXTFLEX Rapid DNA-Seq Kit (Bioo Scientific, Austin, TX, USA). Full-length sequencing primer hybridization sites were incorporated through adapter ligation to blunt-ended DNA fragments. Paired-end sequencing was performed on the Illumina NovaSeq 6000 platform (Illumina Inc., San Diego, CA, USA) at Majorbio Bio-Pharm Technology Co., Ltd. (Shanghai, China) using NovaSeq Reagent Kits according to Illumina protocols (www.illumina.com). Sequencing data generated in this project were deposited in the NCBI database under accession number PRJNA1422992.

### 2.3. Sequence quality control and genome assembly

Raw metagenomic data were processed on the Majorbio Cloud Platform (www.majorbio.com). Adapter sequences and low-quality reads were removed using fastp (v0.20.0) (Chen et al. 2018) based on the following criteria: length < 50 bp, average quality score < 20, or containing ambiguous bases. To remove host-derived sequences, reads were aligned to the *Micropterus salmoides* reference genome (GCA_014851395.1) using BWA (v0.7.9a) (Li and Durbin 2009), and mapped reads together with corresponding mate pairs were excluded. The remaining high-quality reads were assembled using MEGAHIT (v1.1.2) (Li et al. 2015), and contigs ≥ 300 bp were retained for downstream analyses.

### 2.4. Gene prediction, taxonomic classification, and functional annotation

Open reading frames (ORFs) were predicted from assembled contigs using MetaGene (Noguchi, Park, and Takagi 2006). ORFs of at least 100 bp were retained and translated into amino acid sequences using the NCBI translation table. A non-redundant gene catalog was generated using CD-HIT (v4.6.1), with sequence identity and alignment coverage thresholds both set to 90% (Fu et al. 2012). High-quality reads were aligned to the non-redundant gene catalog using SOAPaligner (v2.21) (Li et al. 2008) with a minimum identity threshold of 95%, and gene abundance was calculated and normalized to reads per kilobase per million mapped reads (RPKM). Representative sequences from the non-redundant gene catalog were taxonomically annotated by alignment against the NCBI non-redundant (NR) database using Diamond (v0.8.35), with an e-value cutoff of 10^-5^ (Buchfink, Xie, and Huson 2015). Functional annotation was performed by aligning representative sequences against the Kyoto Encyclopedia of Genes and Genomes (KEGG) database using Diamond (v0.8.35), with an e-value cutoff of 10^-5^. Antibiotic resistance genes (ARGs) and virulence factors (VFs) were annotated using Diamond (v0.8.35) against the Comprehensive Antibiotic Resistance Database (CARD) and Virulence Factor Database (VFDB), respectively, with an e-value threshold of 10^-5^. Carbohydrate-active enzymes were identified using hmmscan (http://hmmer.janelia.org/search/hmmscan) against the CAZy database, with the e-value threshold also set to 10^-5^.

### 2.5. Statistical analysis

Microbial abundance and community composition data are presented as mean ± standard deviation (SD) from five biological replicates per group. Alpha diversity was assessed using ACE, Shannon, and Simpson indices calculated with the phyloseq package (v1.40.0). Group differences in alpha diversity were analyzed by one-way analysis of variance (ANOVA), followed by the Tukey-Kramer test. Beta diversity was calculated from Bray-Curtis distance dissimilarities, and principal coordinates analysis (PCoA) was performed using phyloseq (v1.40.0). Linear discriminant analysis (LDA) effect size (LEfSe) was used to identify taxa that differed significantly among groups, with the significance threshold set at *P* < 0.05. Differences in microbial abundance among groups were evaluated using the Kruskal-Wallis test, followed by *post hoc* Dunn’s multiple comparisons with Bonferroni correction.

### 2.6. Ethical Statement

The care and use of fish samples in this study complied with the institutional animal welfare guidelines and policies of Foshan University. All fish sampling procedures were conducted in accordance with the animal ethics requirements of Foshan University, and were approved by the Animal Ethics Committee of Foshan University.

## 3. Results

### 3.1. Overview of metagenomic sequencing

A total of 20 intestinal samples were collected from largemouth bass across four groups. Metagenomic sequencing generated approximately 879.7 million paired-end (PE) raw reads, with an average of 43.99 million reads per sample (Table S1). After quality control and host genome removal, 227.4 million high-quality reads were retained, with an average of 11.37 million reads per sample. Assembly of the clean reads produced 656 367 contigs, from which 962 480 ORFs were predicted (Table S1). Following de-redundancy, 255 747 non-redundant genes were identified with an average length of 609.33 bp (Table S2), the majority of which clustered within the 200–600 bp range (Fig. S1). Taxonomic annotation of the non-redundant gene set against the NCBI non-redundant (NR) database identified 8 174 species, classified into 2 684 genera, 1 053 families, 524 orders, 129 phyla, 10 kingdoms, and four taxonomic categories (Table S3).

### 3.2. Taxonomic composition of gut microbiota

The gut microbiota of largemouth bass spanned four major taxonomic categories across all groups. Bacteria dominated the microbial community, comprising more than 95% of total sequences in each group (IA: 95.30%, CA: 98.47%, PH: 98.16%, PD: 98.49%) (Fig. 2A). Within the non-bacterial fraction, Eukaryota (3.87%) and Archaea (0.15%) were more abundant in the IA group, while viral sequences were relatively more abundant in the PH group (0.85%) (Fig. 2B–E).

**Figure 2.**
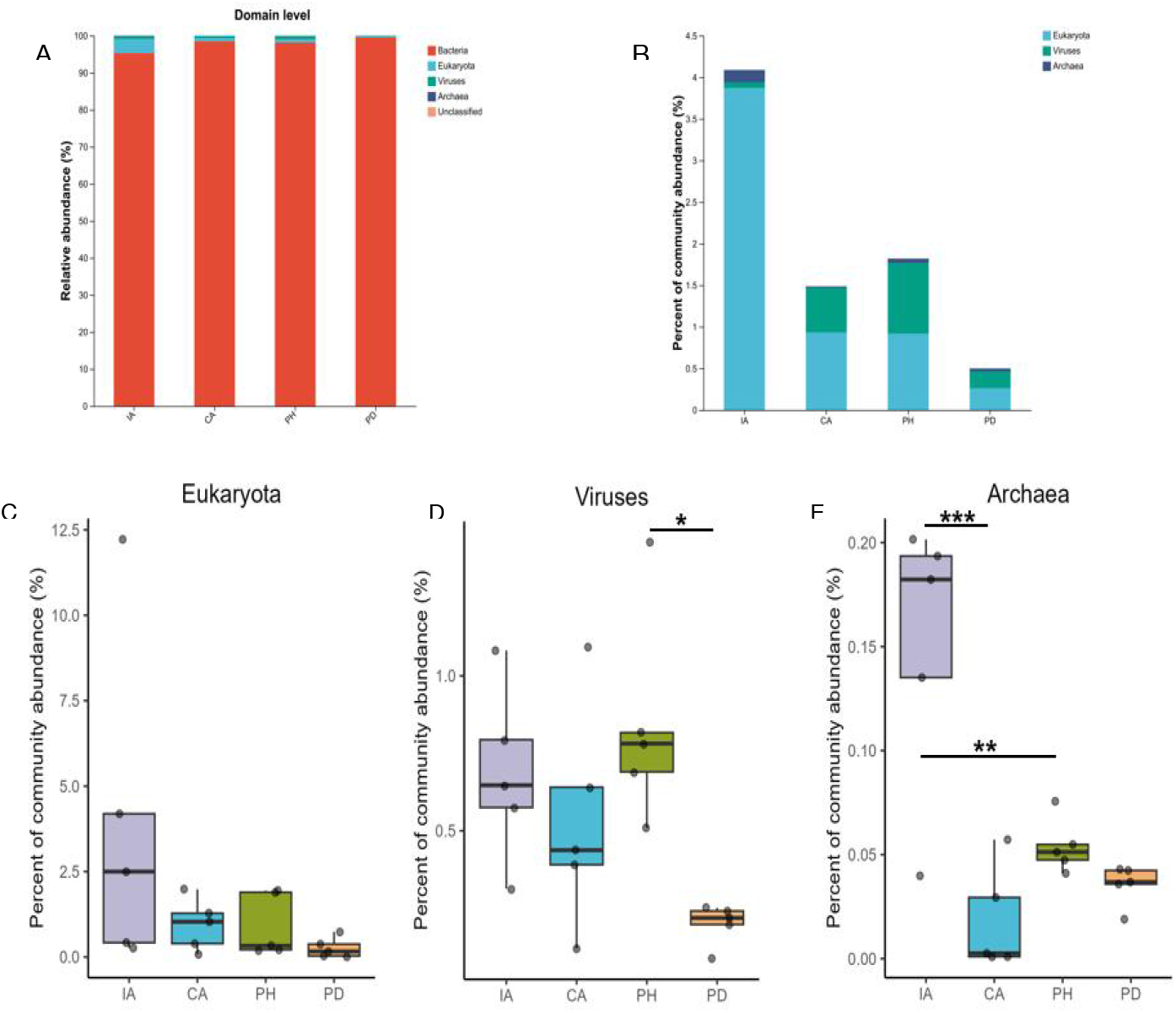
Composition of largemouth bass gut microbiota across taxonomic categories. A. Taxonomic annotation of gut microbial communities. B. Taxonomic annotation of sequences excluding bacteria. C, D, and E. Relative abundances of Eukaryota (C), Viruses (D), and Archaea (E) across four groups.

### 3.3. Influence of aquaculture models on gut microbial abundance and diversity

PCoA revealed distinct clustering among the groups, indicating clear separation in gut microbial community composition across aquaculture models (Fig. 3A). UpSet analysis demonstrated that 510 microbial genera were shared among the IA, CA, and PH groups, with IA fish containing 1 332 unique genera, significantly more than found in the CA (144) and PH fish (47) (Fig. 3B). Assessment of alpha diversity showed that the ACE index, reflecting species richness, was significantly higher in the IA group than in the CA and PH groups (*P* < 0.05) (Fig. 3C). Similarly, the Shannon index, representing microbial diversity, was elevated in the IA group, while the Simpson index did not differ significantly among the three groups (Fig. 3D, E). These results indicate that the IA system was associated with higher gut microbial richness and diversity than the CA and PH systems, potentially contributing to a more stable intestinal microbial ecosystem.

**Figure 3.**
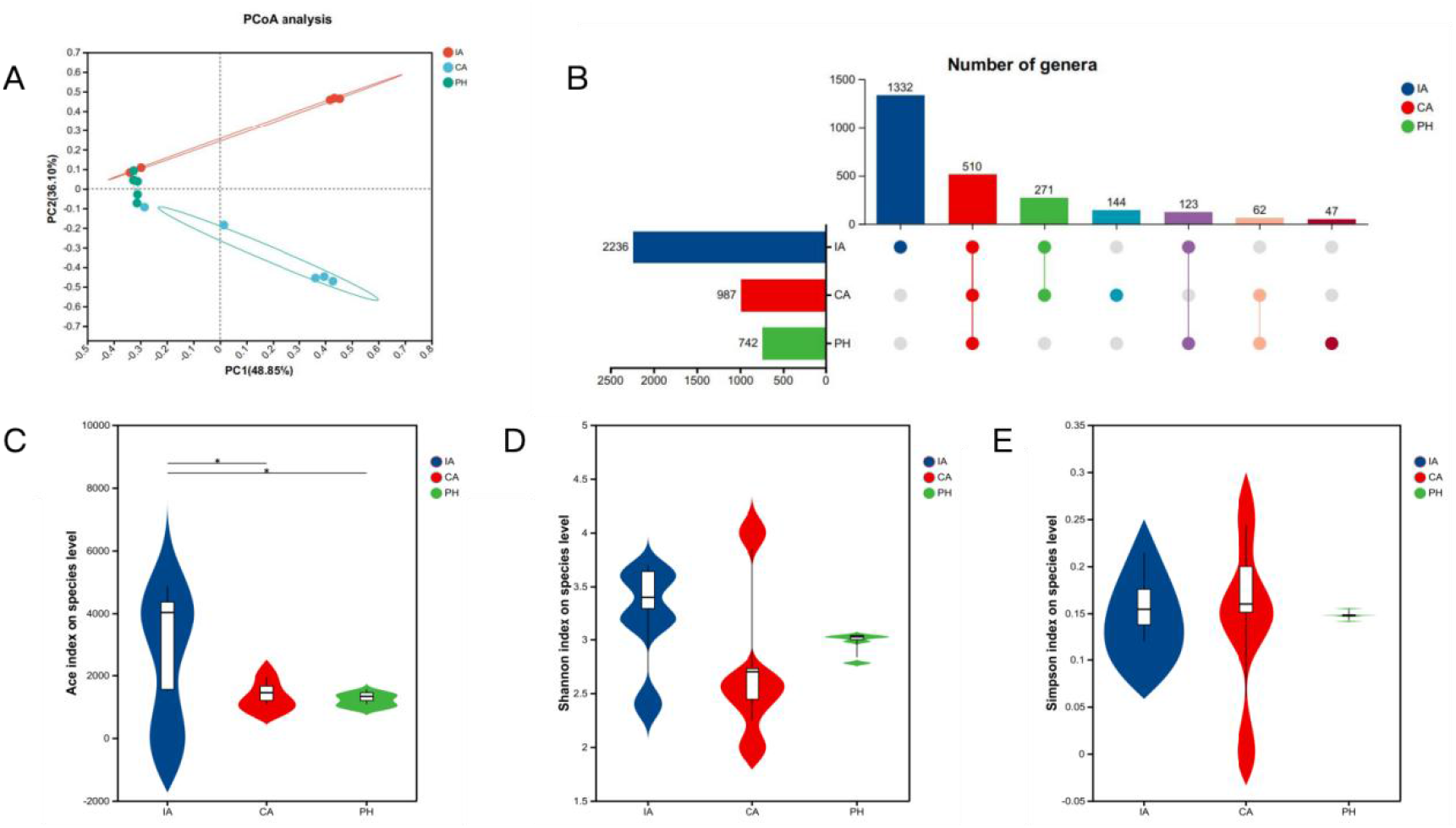
Richness and diversity of intestinal microbiota in largemouth bass across aquaculture models. A. PCoA of the gut microbiota composition. B. UpSet diagram showing shared and unique gut microbiota genera. C. ACE index showing richness of gut microbiota, with higher values indicating greater richness. D–E. Shannon and Simpson indices showing diversity of gut microbiota. CA: pond captive mode; IA: industrial aquaculture model; PH: ordinary pond culture mode (healthy fish). * *P* < 0.05.

### 3.4. Aquaculture models shape the gut microbiome composition of largemouth bass

To evaluate the impact of rearing conditions on gut bacterial composition, comparative analyses were conducted across aquaculture environments. At the phylum level, Fusobacteria and Firmicutes were dominant in all groups (Fig. 4A), with Fusobacteria most abundant in the PH group (71.82%) and Firmicutes highest in the IA group (59.04%). Proteobacteria showed markedly elevated abundance in the CA group (50.50%) compared to the IA and PH groups.

**Figure 4.**
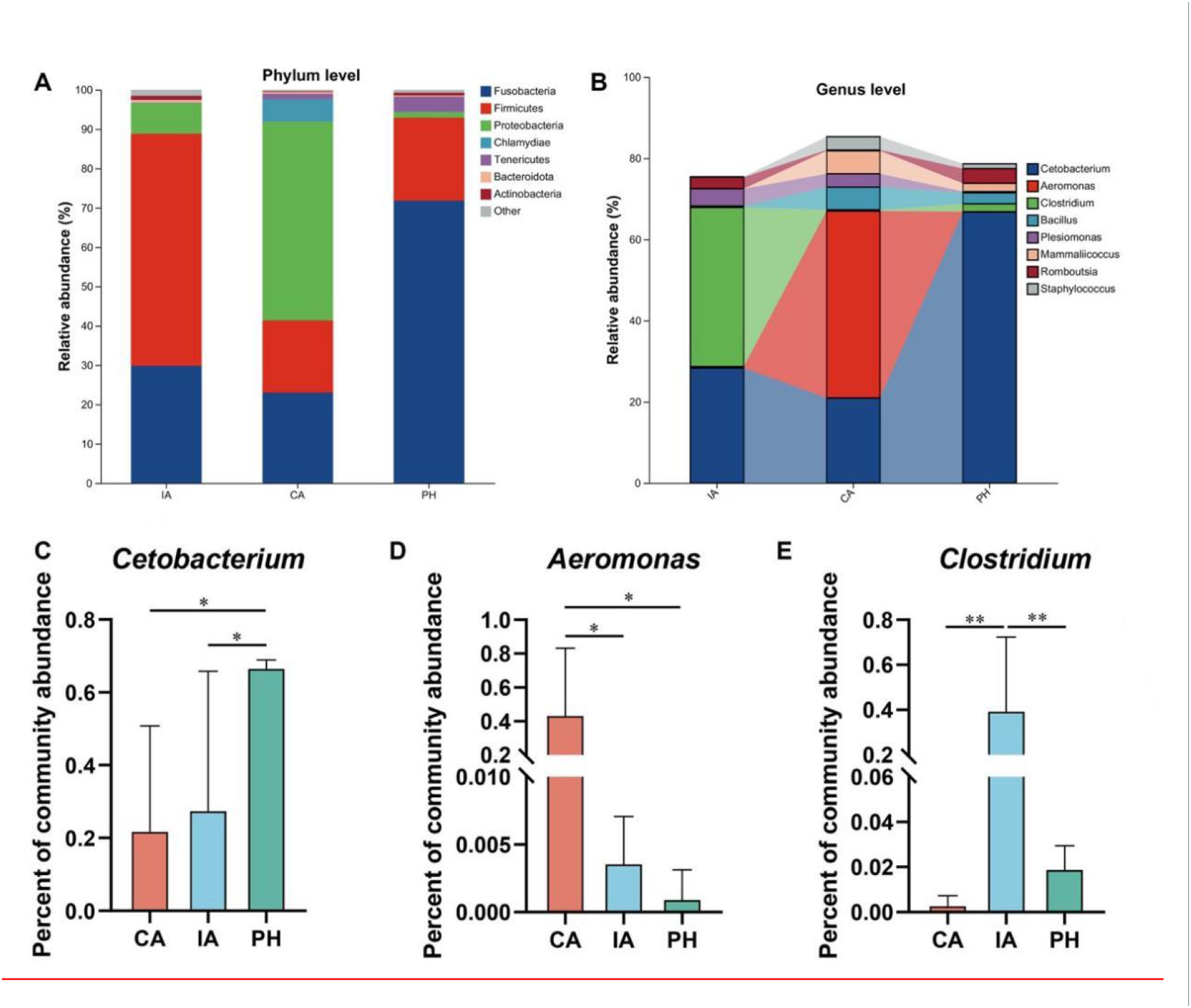
Comparative analysis of gut bacterial composition in largemouth bass across aquaculture environments. A. Phylum-level taxonomic annotation of largemouth bass gut bacteria. B. Genus-level taxonomic annotation of largemouth bass gut bacteria, highlighting *Cetobacterium*, *Aeromonas*, and *Clostridium*. C–E. Relative abundance of *Cetobacterium* (C), *Aeromonas* (D), and *Clostridium* (E) across groups. CA: pond captive mode; IA: industrial aquaculture model; PH: ordinary pond culture mode (healthy fish). * *P* < 0.05 and ** *P* < 0.01.

At the genus level, *Cetobacterium* was prevalent across all groups (Fig. 4B), but was significantly more abundant in the PH group (66.73%) than in the CA (20.94%) and IA (28.31%) groups (*P* < 0.05) (Fig. 4C). In contrast, *Aeromonas* was highly enriched in the CA group (46.05%) but nearly absent in the IA (0.36%) and PH (0.099%) groups (*P* < 0.05) (Fig. 4D). *Clostridium* exhibited its highest abundance in the IA group (39.21%), significantly exceeding its levels in the CA (0.30%) and PH (1.94%) groups (*P* < 0.01) (Fig. 4E).

At the species level, a broad diversity of taxa was annotated in each group (Fig. S2A). Notably, *Aeromonas veronii* and *A. hydrophila* were significantly enriched in the CA group compared to the IA and PH groups (Fig. S2B, C), suggesting a heightened risk of disease associated with this culture model. In contrast, *Clostridium thermobutyricum* and unclassified *Cetobacterium* were dominant in the IA and PH groups, respectively (Fig. S2D). Collectively, these findings indicate that aquaculture environments exert a strong influence on gut bacterial communities in largemouth bass.

### 3.5. Structural differences in intestinal bacterial communities between healthy and diseased fish in pond culture

To investigate taxonomic differences associated with health status, the gut microbiota of healthy and diseased largemouth bass reared in ordinary pond culture was analyzed using metagenomic data. PCoA based on Bray-Curtis distances revealed significant differences in beta diversity between the two groups (Fig. 5A). Similarly, the ACE index showed significantly higher microbial richness in the PD group compared to the PH group (*P* < 0.05) (Fig. 5B). However, no significant differences were observed in alpha diversity based on Simpson (*P* = 0.50) and Shannon indices (*P* = 0.21) (Fig. 5C, D). At the phylum level, the dominant bacterial taxa in both groups were Fusobacteria, Firmicutes, Proteobacteria, Tenericutes, and Bacteroidota (Fig. 5E).

**Figure 5.**
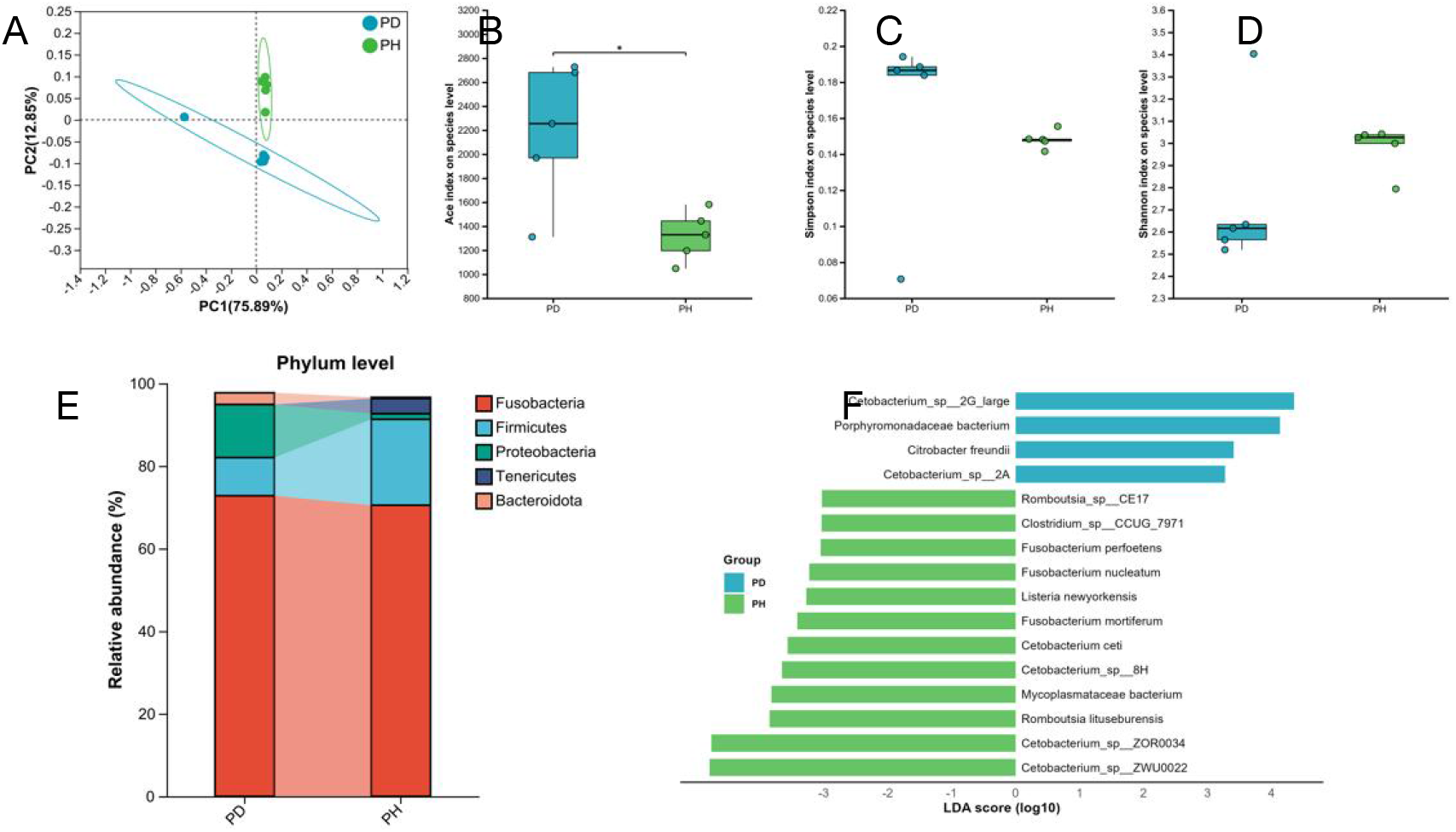
Comparative analysis of gut bacterial communities between healthy and diseased largemouth bass. A. PCoA based on Bray-Curtis distances. B–D. Alpha diversity comparisons based on ACE, Simpson, and Shannon indices. E. Phylum-level relative abundance of gut microbiota visualized using a stacked bar plot. F. LEfSe analysis identifying differentially abundant taxa; columns represent biomarkers with an LDA score > 3.

LEfSe identified several taxa with significantly different relative abundance between the two groups (Fig. 5F). The PD group was enriched in taxa such as *Cetobacterium* sp., *Porphyromonadaceae bacterium*, and *Citrobacter freundii*, the latter being a potential pathogen associated with intestinal dysbiosis. In contrast, commensal species such as *Romboutsia lituseburensis* were more abundant in the PH group.

### 3.6. Functional profiles of gut microbiota genes vary across aquaculture models

To assess the functional potential of the gut microbiota, predicted genes were annotated using the KEGG database to identify associated metabolic pathways. Among the top 10 most enriched pathways, five exhibited significant differences in relative abundance across the four aquaculture groups (Fig. 6A). Functional comparisons between healthy and diseased fish under ordinary pond culture further revealed distinct enrichment patterns (Fig. 6B). Pathways related to purine metabolism, alanine, aspartate, and glutamate metabolism, and fatty acid metabolism were enriched in the PD group, while O-antigen nucleotide sugar biosynthesis was enriched in the PH group. These differences suggest disease-associated shifts in the predicted functional potential of the gut microbiota.

**Figure 6.**
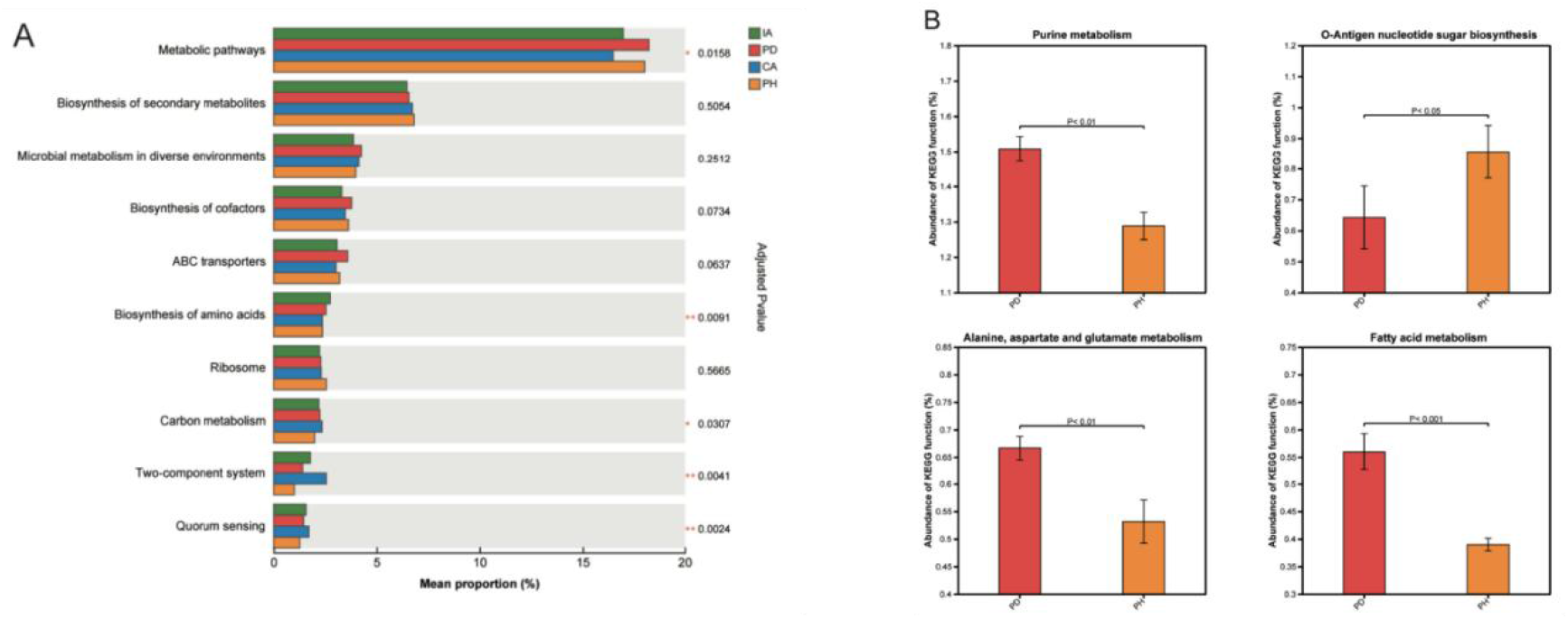
KEGG pathway analysis of gut microbiota genes. A. Five of the top 10 pathways differed significantly across the four aquaculture groups. B. Comparison of metabolic pathway enrichment between PD and PH groups.

### 3.7. Virulence factors, carbohydrate-active enzymes, and antibiotic resistance genes

Metagenomic analysis revealed multiple types of virulence factors (VFs) in the gut microbiota of largemouth bass, spanning categories such as toxins, magnesium uptake systems, antiphagocytosis, serum resistance, secretion systems, adherence, and iron uptake systems, based on level 1 functional classification. LEfSe identified group-specific VF signatures (LDA > 3.5; *P* < 0.05; Fig. 7A). The CA group was enriched for secretion system-associated VFs, including VF0479, VF0344, and SS194. Notably, the PD group showed enrichment of capsule-associated factors, including VF0543 and VF0274, while the PH group showed higher levels of adherence-associated factors such as VF0326, indicating distinct virulence profiles associated with rearing conditions and physiological state.

**Figure 7.**
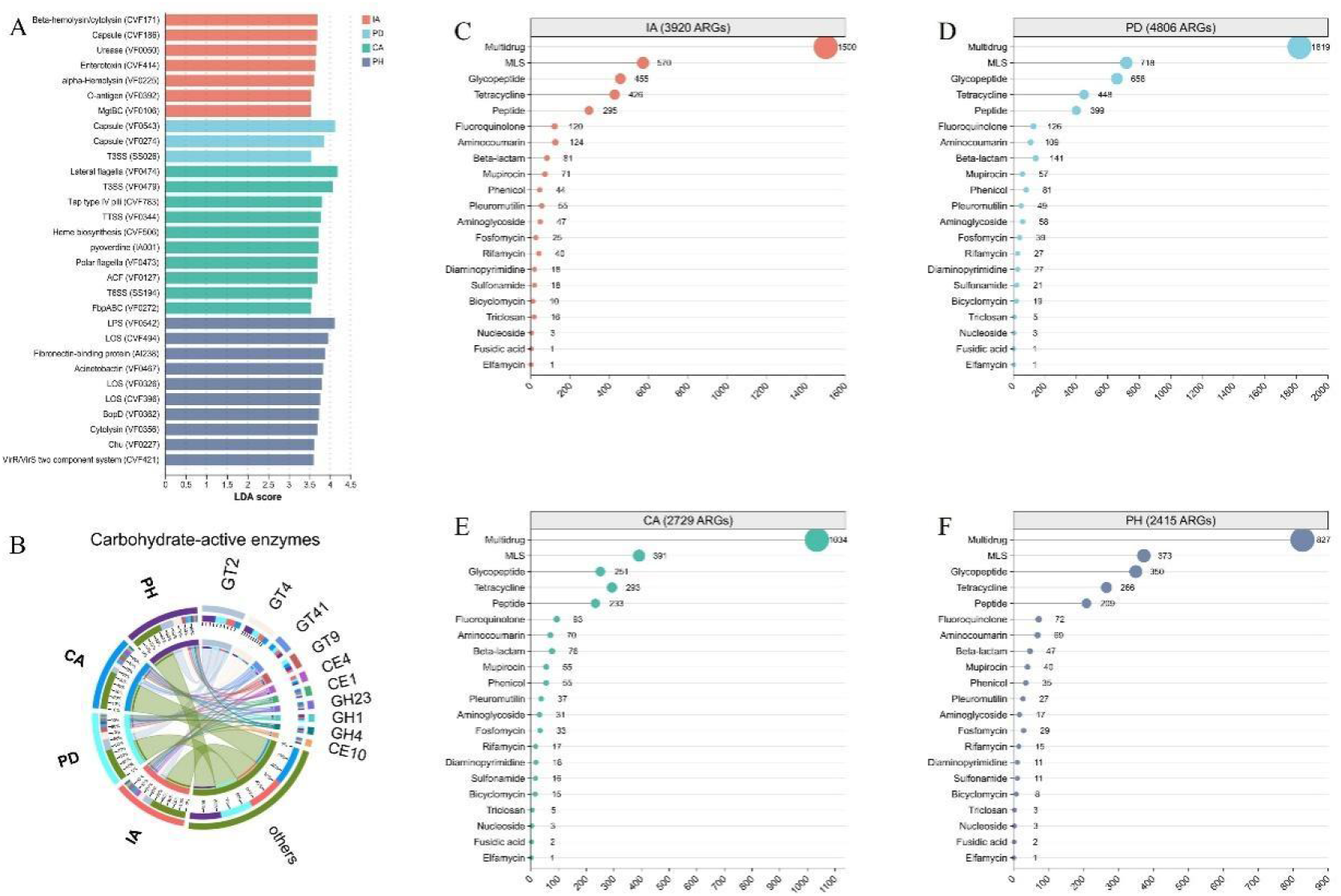
Gene profiles of the gut microbiota under different aquaculture models. A. Identification of VF signatures in each group based on LEfSe analysis (LDA > 3.5; *P* < 0.05). B. Circos plot showing the distribution of carbohydrate-active enzymes across groups. C–F. Bubble charts depicting the abundance of antibiotic classes under different aquaculture models.

To further evaluate functional differentiation, the composition and distribution of carbohydrate-active enzymes in the gut microbiota were assessed across groups. Circos visualization revealed a diverse repertoire of carbohydrate-active enzymes across groups, including glycosyltransferases, carbohydrate esterases, and glycoside hydrolases (Fig. 7B). These enzyme profiles indicate broad differences in carbohydrate-processing potential among groups.

Comparative analysis against a comprehensive ARG database identified thousands of ARGs, distributed among 21 antibiotic classes (Fig. 7C–F). The number of detected ARGs varied across aquaculture models, with the highest count in the PD group (4 806), followed by IA (3 920), CA (2 729), and PH (2 415). Across all groups, multidrug resistance and macrolide-lincosamide-streptogramin (MLS) resistance genes were most prevalent. Notably, MLS-type ARGs were most abundant in the IA group and lowest in the CA and PH groups. Several ARG classes displayed aquaculture-specific enrichment patterns, such as significantly higher multidrug ARG abundance in the IA group, potentially associated with differences in antibiotic exposure or farming practices among systems. The ARG profile in the PH group also differed from those in the CA and IA groups, potentially indicating functional differences in the gut microbiota of healthy fish. Notably, the PD and PH groups differed significantly in ARG number (Fig. 7D, F), indicating marked disease-associated shifts in gut resistome diversity and predicted microbial function.

## 4. Discussion

Gut microbial communities contribute to host physiology in fish by supporting nutrient processing, intestinal immune regulation, and resistance to opportunistic infection (Ni et al. 2014; Larsen, Mohammed, and Arias 2014; Li et al. 2014; Yan et al. 2016; Dulski, Zakęś, and Ciesielski 2018; Burgos, Cai, and Arias 2024). Water quality, feeding management, and stocking density can alter intestinal microbial structure by changing nutrient availability, physicochemical conditions, and exposure to environmental microbes (Yan et al. 2017; Zhang et al. 2025; Groves et al. 2023). In this study, largemouth bass reared under different aquaculture regimes showed pronounced differences in gut microbial diversity and taxonomic composition. Notably, IA fish exhibited higher microbial richness and diversity, likely reflecting the tightly controlled and relatively stable physicochemical parameters of this rearing environment. In contrast, CA fish harbored elevated levels of opportunistic pathogens, such as *A. veronii* and *A. hydrophila,* organisms frequently implicated in disease outbreaks. The semi-open nature of pond enclosures, which exposes fish to fluctuating environmental inputs, can create favorable conditions for pathogen proliferation and transmission. Enrichment of pathogenic taxa in CA fish highlights the increased vulnerability of open systems to microbial imbalance and underscores the need for improved biosecurity measures (Zhang et al. 2014). These findings are consistent with previous reports demonstrating increased pathogen burden in fish reared under open aquaculture conditions (Lafferty et al. 2015), reinforcing the ecological importance of environmental management in disease prevention.

Although the healthy (PH) and diseased (PD) groups were both derived from pond aquaculture, their gut microbiota displayed markedly different taxonomic profiles. This divergence suggests that disease status profoundly impacts the structure of the gut microbial community. Pathophysiological changes during infection likely disrupt the intestinal microenvironment, promoting a shift toward dysbiosis characterized by reduced beneficial taxa and increased colonization by harmful species. This imbalance may further weaken host immunity, facilitate pathogen persistence, and exacerbate disease progression. The observed microbiota shifts associated with disease status underscore the value of microbial monitoring. Timely profiling of gut microbiota during culture operations may enable early detection of dysbiosis and support interventions to mitigate disease risk (El-Son et al. 2025).

At the phylum level, Fusobacteria and Firmicutes were consistently dominant across all groups, although their relative abundances displayed notable variation. The PH group exhibited the highest abundance of Fusobacteria, while Firmicutes predominated in the IA group. Members of Fusobacteria have been implicated in protein degradation and short-chain fatty acid production, processes that facilitate nutrient assimilation in carnivorous fish. Their elevated abundance in the PH group may contribute to enhanced digestive efficiency and metabolic performance. In contrast, the prominence of Firmicutes in the IA group may be linked to carbohydrate fermentation, energy metabolism, and host immune modulation. The higher microbial diversity observed in the IA group may facilitate interactions between Firmicutes and other taxa, thereby supporting more efficient nutrient utilization and contributing to host immunocompetence (Sharmin et al. 2013; Zhang et al. 2024; Gallet et al. 2023). At the genus level, *Cetobacterium* was present across all groups, with the highest abundance in the PH group. *Cetobacterium* is known to synthesize key micronutrients such as vitamin B12 and may play a critical role in maintaining mucosal integrity, promoting nutrient absorption, and supporting systemic physiological functions (Qi et al. 2023; Wang et al. 2021; Tsuchiya, Sakata, and Sugita 2008). Its enrichment in healthy fish suggests a potential association with intestinal homeostasis and favorable growth-related physiology. Conversely, enrichment of the potential pathogen *Citrobacter freundii* in the PD group underscores the association between microbial dysbiosis and disease. As an opportunistic pathogen, *C. freundii* is capable of disrupting the intestinal mucosal barrier, producing enterotoxins, and triggering inflammatory responses through competitive displacement of commensals and resource monopolization. These pathogenic mechanisms may contribute to intestinal damage and increased disease susceptibility in infected fish.

Functional prediction analysis revealed an association between physiological status and the metabolic profile of the gut microbiome. In the PD group, pathways related to purine and fatty acid metabolism were significantly enriched, suggesting changes in predicted energy-related and immune-associated functions under disease conditions. Purine metabolism is involved in cellular energy supply and signal transduction, and its enrichment in diseased fish may reflect increased metabolic demand during pathogen invasion and tissue repair. Fatty acid metabolism not only provides energy but also contributes to cell membrane composition and lipid-mediated signaling. Enrichment of fatty acid metabolism in the PD group suggests regulation of membrane composition, energy generation, and lipid-based signaling to support immune cell activation and intercellular communication during infection. In contrast, the PH group exhibited enrichment of pathways involved in O-antigen nucleotide sugar biosynthesis, reflecting enhanced intestinal barrier function and immune response in healthy fish. O-antigens, key components of bacterial outer membranes, are involved in microbial adhesion, immune recognition, and modulation of host-microbe interactions (Zhao et al. 2022). These patterns suggest that microbial metabolic functions differ between healthy and diseased fish and may be linked to intestinal immune status.

This study identified distinct gut microbiome profiles across major largemouth bass aquaculture systems, providing reference data to support microbiome-based health management in aquaculture. Future large-scale controlled trials should determine how specific environmental and husbandry interventions affect fish growth, immune status, product quality, and microbiome stability. Such work will help define practical strategies that integrate microbial regulation into sustainable aquaculture management.

## Supporting information

Supplementary Table S1. Sequencing data and assembly statistics

Supplementary Table S2. Gene catalog redundancy statistics

Supplementary Table S3. Species taxonomic annotations against NR database

## Acknowledgements

This research was funded by the APRC-CityU New Research Initiatives/Infrastructure Support (9610574) and the SIRG-CityU Strategic Interdisciplinary Research Grant (7020090).

## Data availability

All sequencing data and genome assemblies have been deposited in NCBI BioProject under accession number PRJNA1422992.

