## Supplementary Table S1. Sequencing data and assembly statistics for "Meta-analysis of gut microbiome in largemouth bass (Micropterus salmoides) under different aquaculture systems"

| | Raw data | | After quality control | | | | Remove host sequences | | | | Assemble | | | | | | Gene prediction | | | | |
|---|---|---|---|---|---|---|---|---|---|---|---|---|---|---|---|---|---|---|---|---|---|
| Samples | Raw reads | Raw base (bp) | Clean reads | Clean base (bp) | Percent in raw reads(%) | Percent in raw bases(%) | Optimized reads | Optimized bases (bp) | Percent in raw reads (%) | Percent in raw bases (%) | Contigs | Contigs bases(bp) | N50(bp) | N90(bp) | Max(bp) | Min(bp) | ORFs | Total Length(bp) | Average Length(bp) | Max(bp) | Min(bp) |
| CA_1 | 42150738 | 6364761438 | 41555478 | 6256959054 | 98.587782733484 | 98.30626198562 | 26377896 | 3978574554 | 62.579914970884 | 62.509405776726 | 50875 | 50025066 | 1535 | 382 | 137026 | 300 | 79785 | 43515813 | 545.41 | 15792 | 102 |
| CA_2 | 41621064 | 6284780664 | 40896324 | 6144153363 | 98.258718229789 | 97.762415133983 | 1318740 | 198547757 | 3.1684437476178 | 3.1591835517396 | 14771 | 12524012 | 950 | 383 | 840204 | 300 | 20950 | 10680636 | 509.82 | 7227 | 102 |
| CA_3 | 43397840 | 6553073840 | 42780322 | 6412810508 | 98.577076647133 | 97.859579558774 | 12642018 | 1904930121 | 29.130523546794 | 29.069260739476 | 25797 | 33471950 | 3030 | 437 | 769643 | 300 | 48114 | 28988322 | 602.49 | 16299 | 102 |
| CA_4 | 45939668 | 6936889868 | 45142026 | 6769829629 | 98.263718405627 | 97.591712681347 | 6743112 | 1014069576 | 14.678190534594 | 14.618504766494 | 45396 | 39721141 | 1138 | 370 | 126371 | 300 | 66157 | 34139652 | 516.04 | 11949 | 102 |
| CA_5 | 42615584 | 6434953184 | 41943062 | 6304024476 | 98.421887166911 | 97.96535104054 | 2421926 | 365064394 | 5.6831932656373 | 5.6731476292276 | 36960 | 24119418 | 692 | 344 | 25224 | 300 | 46996 | 20602905 | 438.4 | 4650 | 102 |
| IA_1 | 46311598 | 6993051298 | 45162604 | 6757262908 | 97.518992974503 | 96.628247385123 | 15239688 | 2286076068 | 32.906849813302 | 32.690680656866 | 81236 | 63924986 | 771 | 362 | 937120 | 300 | 86587 | 44240613 | 510.94 | 15111 | 102 |
| IA_2 | 42062222 | 6351395522 | 41086828 | 6140749206 | 97.681068774731 | 96.683464047069 | 9887468 | 1480910211 | 23.506765762398 | 23.316296487448 | 99045 | 66986465 | 635 | 357 | 937089 | 300 | 75580 | 34536936 | 456.96 | 15111 | 102 |
| IA_4 | 43843812 | 6620415612 | 42855774 | 6403335320 | 97.746459637223 | 96.721047367381 | 6312352 | 948548496 | 14.397361251344 | 14.327627623267 | 16301 | 19293845 | 2008 | 441 | 143190 | 300 | 28409 | 16757502 | 589.87 | 15723 | 102 |
| IA_5 | 41482346 | 6263834246 | 40571410 | 6044366997 | 97.804039337602 | 96.496279429167 | 4342674 | 651625363 | 10.468728070491 | 10.402979028638 | 15473 | 19116549 | 3202 | 413 | 219019 | 300 | 27613 | 16734003 | 606.02 | 15723 | 102 |
| IA_6 | 45101856 | 6810380256 | 43853580 | 6555271584 | 97.232317889534 | 96.254119999023 | 9621012 | 1442430467 | 21.331742977495 | 21.179881486488 | 32173 | 35169136 | 2673 | 389 | 937120 | 300 | 50317 | 28792836 | 572.23 | 15111 | 102 |
| PD_4 | 44636838 | 6740162538 | 43531306 | 6523852059 | 97.523274386058 | 96.790723105259 | 20649944 | 3102381840 | 46.26211202505 | 46.028294162184 | 48082 | 61005185 | 2477 | 444 | 379841 | 300 | 85089 | 53248245 | 625.79 | 18414 | 102 |
| PD_7 | 45492500 | 6869367500 | 44420122 | 6631023248 | 97.642736714843 | 96.530331911926 | 3639274 | 546808355 | 7.9997230312689 | 7.9600975635675 | 7288 | 12109351 | 5388 | 529 | 248258 | 300 | 15357 | 10319580 | 671.98 | 15795 | 102 |
| PD_8 | 48587484 | 7336710084 | 47543324 | 7042654664 | 97.850969191984 | 95.991998911865 | 8025082 | 1205686922 | 16.516767980824 | 16.433618177572 | 23467 | 26443054 | 1783 | 433 | 248258 | 300 | 40549 | 23110686 | 569.94 | 15795 | 102 |
| PD_9 | 44644526 | 6741323426 | 43654344 | 6530372318 | 97.782075231351 | 96.870776038034 | 21755328 | 3266335090 | 48.730113071421 | 48.452431126541 | 42998 | 47674304 | 1617 | 441 | 275137 | 300 | 73418 | 41652069 | 567.33 | 15795 | 102 |
| PD_10 | 46531908 | 7026318108 | 45449408 | 6823732413 | 97.673639344426 | 97.116758850281 | 33851370 | 5084773035 | 72.74872545523 | 72.367532423711 | 47783 | 58140828 | 1912 | 465 | 275137 | 300 | 86811 | 50871201 | 586 | 15795 | 102 |
| PH_1 | 41248020 | 6228451020 | 40692058 | 6121619314 | 98.652148636468 | 98.284778901577 | 10696830 | 1613464014 | 25.932953872695 | 25.904739538274 | 19655 | 24830268 | 2114 | 477 | 113641 | 300 | 35207 | 21415650 | 608.28 | 11244 | 102 |
| PH_2 | 43298298 | 6538042998 | 42657420 | 6417404279 | 98.519854059852 | 98.154819124975 | 14904196 | 2247666394 | 34.422129017635 | 34.378274885735 | 15279 | 27717003 | 8249 | 527 | 518932 | 300 | 35884 | 24110403 | 671.9 | 15687 | 102 |
| PH_3 | 42612148 | 6434434348 | 42002138 | 6316909117 | 98.56845986736 | 98.173495529774 | 15132488 | 2282928580 | 35.512145503672 | 35.479864375485 | 15766 | 24158463 | 3232 | 507 | 242477 | 300 | 31856 | 20934813 | 657.17 | 11646 | 102 |
| PH_4 | 43045386 | 6499853286 | 42411220 | 6379696593 | 98.526750346715 | 98.151393766705 | 2108574 | 317795233 | 4.8984901657056 | 4.889267788313 | 9262 | 9818782 | 1758 | 402 | 72207 | 300 | 14289 | 8284596 | 579.79 | 10548 | 102 |
| PH_5 | 45089902 | 6808575202 | 44298996 | 6659445861 | 98.245935420308 | 97.809683574381 | 1725310 | 260072423 | 3.8263777996235 | 3.8197774906504 | 8760 | 8891259 | 1311 | 406 | 156182 | 300 | 13512 | 7243875 | 536.11 | 11547 | 102 |
| In total | =SUM(B3:B22) | =SUM(C3:C22) | =SUM(D3:D22) | =SUM(E3:E22) | - | - | =SUM(H3:H22) | =SUM(I3:I22) | - | - | =SUM(L3:L22) | =SUM(M3:M22) | - | - | - | - | =SUM(R3:R22) | =SUM(S3:S22) | - | - | - |
| Mean | =AVERAGE(B3:B22) | =AVERAGE(C3:C22) | =AVERAGE(D3:D22) | =AVERAGE(E3:E22) | =AVERAGE(F3:F22) | =AVERAGE(G3:G22) | =AVERAGE(H3:H22) | =AVERAGE(I3:I22) | =AVERAGE(J3:J22) | =AVERAGE(K3:K22) | =AVERAGE(L3:L22) | =AVERAGE(M3:M22) | =AVERAGE(N3:N22) | =AVERAGE(O3:O22) | =AVERAGE(P3:P22) | =AVERAGE(Q3:Q22) | =AVERAGE(R3:R22) | =AVERAGE(S3:S22) | =AVERAGE(T3:T22) | =AVERAGE(U3:U22) | =AVERAGE(V3:V22) |
