## Supplementary Table S2. Gene catalog redundancy statistics for "Meta-analysis of gut microbiome in largemouth bass (Micropterus salmoides) under different aquaculture systems"

### Sheet1

| Before redundancy | | | After redundancy | | |
|---|---|---|---|---|---|
| **Genes** | **Total length (bp)** | **Average length (bp)** | **Catalog genes** | **Catalog total length (bp)** | **Catalog average length (bp)** |
| 962480 | 540180336 | 561.24 | 255747 | 155833125 | 609.33 |
