## Supplementary materials for "Meta-analysis of gut microbiome in largemouth bass (Micropterus salmoides) under different aquaculture systems"

### Supplementary Table S4 - Table S1 (Assembly statistics)

| | Raw data | After quality control | | Remove host sequences | | Assemble | | | | | | Gene prediction | | | | |
|---|---|---|---|---|---|---|---|---|---|---|---|---|---|---|---|---|
| Samples | Raw reads | Clean reads | Percent in raw reads(%) | Optimized reads | Percent in raw reads (%) | Contigs | Contigs bases(bp) | N50(bp) | N90(bp) | Max(bp) | Min(bp) | ORFs | Total Length(bp) | Average Length(bp) | Max(bp) | Min(bp) |
| CA_1 | 42150738 | 41555478 | 98.587782733484 | 26377896 | 62.579914970884 | 50875 | 50025066 | 1535 | 382 | 137026 | 300 | 79785 | 43515813 | 545.41 | 15792 | 102 |
| CA_2 | 41621064 | 40896324 | 98.258718229789 | 1318740 | 3.1684437476178 | 14771 | 12524012 | 950 | 383 | 840204 | 300 | 20950 | 10680636 | 509.82 | 7227 | 102 |
| CA_3 | 43397840 | 42780322 | 98.577076647133 | 12642018 | 29.130523546794 | 25797 | 33471950 | 3030 | 437 | 769643 | 300 | 48114 | 28988322 | 602.49 | 16299 | 102 |
| CA_4 | 45939668 | 45142026 | 98.263718405627 | 6743112 | 14.678190534594 | 45396 | 39721141 | 1138 | 370 | 126371 | 300 | 66157 | 34139652 | 516.04 | 11949 | 102 |
| CA_5 | 42615584 | 41943062 | 98.421887166911 | 2421926 | 5.6831932656373 | 36960 | 24119418 | 692 | 344 | 25224 | 300 | 46996 | 20602905 | 438.4 | 4650 | 102 |
| IA_1 | 46311598 | 45162604 | 97.518992974503 | 15239688 | 32.906849813302 | 81236 | 63924986 | 771 | 362 | 937120 | 300 | 86587 | 44240613 | 510.94 | 15111 | 102 |
| IA_2 | 42062222 | 41086828 | 97.681068774731 | 9887468 | 23.506765762398 | 99045 | 66986465 | 635 | 357 | 937089 | 300 | 75580 | 34536936 | 456.96 | 15111 | 102 |
| IA_4 | 43843812 | 42855774 | 97.746459637223 | 6312352 | 14.397361251344 | 16301 | 19293845 | 2008 | 441 | 143190 | 300 | 28409 | 16757502 | 589.87 | 15723 | 102 |
| IA_5 | 41482346 | 40571410 | 97.804039337602 | 4342674 | 10.468728070491 | 15473 | 19116549 | 3202 | 413 | 219019 | 300 | 27613 | 16734003 | 606.02 | 15723 | 102 |
| IA_6 | 45101856 | 43853580 | 97.232317889534 | 9621012 | 21.331742977495 | 32173 | 35169136 | 2673 | 389 | 937120 | 300 | 50317 | 28792836 | 572.23 | 15111 | 102 |
| PD_4 | 44636838 | 43531306 | 97.523274386058 | 20649944 | 46.26211202505 | 48082 | 61005185 | 2477 | 444 | 379841 | 300 | 85089 | 53248245 | 625.79 | 18414 | 102 |
| PD_7 | 45492500 | 44420122 | 97.642736714843 | 3639274 | 7.9997230312689 | 7288 | 12109351 | 5388 | 529 | 248258 | 300 | 15357 | 10319580 | 671.98 | 15795 | 102 |
| PD_8 | 48587484 | 47543324 | 97.850969191984 | 8025082 | 16.516767980824 | 23467 | 26443054 | 1783 | 433 | 248258 | 300 | 40549 | 23110686 | 569.94 | 15795 | 102 |
| PD_9 | 44644526 | 43654344 | 97.782075231351 | 21755328 | 48.730113071421 | 42998 | 47674304 | 1617 | 441 | 275137 | 300 | 73418 | 41652069 | 567.33 | 15795 | 102 |
| PD_10 | 46531908 | 45449408 | 97.673639344426 | 33851370 | 72.74872545523 | 47783 | 58140828 | 1912 | 465 | 275137 | 300 | 86811 | 50871201 | 586 | 15795 | 102 |
| PH_1 | 41248020 | 40692058 | 98.652148636468 | 10696830 | 25.932953872695 | 19655 | 24830268 | 2114 | 477 | 113641 | 300 | 35207 | 21415650 | 608.28 | 11244 | 102 |
| PH_2 | 43298298 | 42657420 | 98.519854059852 | 14904196 | 34.422129017635 | 15279 | 27717003 | 8249 | 527 | 518932 | 300 | 35884 | 24110403 | 671.9 | 15687 | 102 |
| PH_3 | 42612148 | 42002138 | 98.56845986736 | 15132488 | 35.512145503672 | 15766 | 24158463 | 3232 | 507 | 242477 | 300 | 31856 | 20934813 | 657.17 | 11646 | 102 |
| PH_4 | 43045386 | 42411220 | 98.526750346715 | 2108574 | 4.8984901657056 | 9262 | 9818782 | 1758 | 402 | 72207 | 300 | 14289 | 8284596 | 579.79 | 10548 | 102 |
| PH_5 | 45089902 | 44298996 | 98.245935420308 | 1725310 | 3.8263777996235 | 8760 | 8891259 | 1311 | 406 | 156182 | 300 | 13512 | 7243875 | 536.11 | 11547 | 102 |
| In total | =SUM(B3:B22) | =SUM(C3:C22) | - | =SUM(E3:E22) | - | =SUM(G3:G22) | =SUM(H3:H22) | - | - | - | - | =SUM(M3:M22) | =SUM(N3:N22) | - | - | - |
| Mean | =AVERAGE(B3:B22) | =AVERAGE(C3:C22) | =AVERAGE(D3:D22) | =AVERAGE(E3:E22) | =AVERAGE(F3:F22) | =AVERAGE(G3:G22) | =AVERAGE(H3:H22) | =AVERAGE(I3:I22) | =AVERAGE(J3:J22) | =AVERAGE(K3:K22) | =AVERAGE(L3:L22) | =AVERAGE(M3:M22) | =AVERAGE(N3:N22) | =AVERAGE(O3:O22) | =AVERAGE(P3:P22) | =AVERAGE(Q3:Q22) |

### Supplementary Table S4 - Table S2 (Gene redundancy)

| Before redundancy | | | After redundancy | | |
|---|---|---|---|---|---|
| Genes | Total length (bp) | Average length (bp) | Catalog genes | Catalog total length (bp) | Catalog average length (bp) |
| 962480 | 540180336 | 561.24 | 255747 | 155833125 | 609.33 |
